# Multistage and transmission-blocking targeted antimalarials discovered from the open-source MMV Pandemic Response Box

**DOI:** 10.1101/2020.06.05.133405

**Authors:** Janette Reader, Mariёtte E. van der Watt, Dale Taylor, Claire Le Manach, Nimisha Mittal, Sabine Ottilie, Anjo Theron, Phanankosi Moyo, Erica Erlank, Luisa Nardini, Nelius Venter, Sonja Lauterbach, Belinda Bezuidenhout, Andre Horatscheck, Ashleigh van Heerden, Grant A. Boyle, David Calvo, Dalu Mancama, Theresa L. Coetzer, Elizabeth A. Winzeler, James Duffy, Lizette L. Koekemoer, Gregory Basarab, Kelly Chibale, Lyn-Marié Birkholtz

## Abstract

New chemical matter is needed to target the divergent biology associated with the different life cycle stages of *Plasmodium*. Here, we report the parallel screening of the Medicines for Malaria Venture Pandemic Response Box to identify multistage-active and stage-specific compounds against various life cycle stages of *Plasmodium* parasites (asexual parasites, stage IV/V gametocytes, gametes, oocysts and liver stages) and for endectocidal activity. Hits displayed unique chemotypes and included two multistage-active compounds, 16 asexual-targeted, six with prophylactic potential and ten gametocyte-targeted compounds. Notably, four structurally diverse gametocyte-targeted compounds with potent transmission-blocking activity were identified: the JmjC inhibitor ML324, two azole antifungals including eberconazole, and the antitubercular clinical candidate SQ109. Besides ML324, none of these have previously attributed antiplasmodial activity, emphasizing the success of *de novo* parallel screening against different *Plasmodium* stages to deliver leads with novel modes-of-action. Importantly, the discovery of such transmission-blocking targeted compounds covers a previously unexplored base for delivery of compounds required for malaria elimination strategies.

## INTRODUCTION

Malaria treatment solely relies on drugs that target the parasite but current treatment options have a finite lifespan due to resistance development. Moreover, whilst current antimalarials are curative of asexual blood stage parasitemia and associated malaria symptoms, they cannot all be used prophylactically and typically do not effectively block transmission. This limits their utility in malaria elimination strategies, where the latter dictates that chemotypes should block human-to-mosquito (gametocyte and gametes) and mosquito-to-human (sporozoites and liver schizonts) transmission.

The transmission stages of malaria parasites are seen as parasite population bottlenecks,^1^ with as few as 100 sporozoites able to initiate an infection after migrating to the liver where exoerythrocytic schizogony occurs. The subsequent release of thousands of daughter cells, which in turn infect erythrocytes, initiates the extensive population expansion that occurs during asexual replication. A minor proportion (~1%)^2^ of the proliferating asexual parasites will undergo sexual differentiation to form mature stage V gametocytes, a 10-14 day process in the most virulent parasite *Plasmodium falciparum*. Only ~10^3^ of these falciform-shaped mature gametocytes are taken up by the next feeding mosquito to transform into male and female gametes in the mosquito’s midgut.^3^ Fertilization results in zygote development, and a motile ookinete that passes through the midgut wall forms an oocyst from which sporozoites develop, making the mosquito infectious.

The sporozoite and gametocyte population bottlenecks have been the basis of enticing arguments towards the development of chemotypes able to target them. However, most compounds able to kill asexual parasites are either ineffective in preventing infection (hepatic development) and blocking transmission (gametocytogenesis) or are compromised by resistance development (e.g. antifolates active as prophylactics). Some compounds also have toxicity concerns (e.g. primaquine targeting gametocytes with associated hemolytic toxicity in glucose-6-phosphate dehydrogenase deficient patients). Patients treated with current antimalarials or asymptomatic carries, may have sufficient levels of gametocytes that can be transmitted to the mosquito and sustain the malaria burden. The development of gametocyte-targeted transmission-blocking compounds is therefore essential for a complete strategy directed at eliminating malaria.

Phenotypic screenings of millions of compounds have successfully identified new antimalarial hits to populate the drug discovery pipeline. However, the majority of these screens assessed activity against asexual blood stage parasites as the primary filter, and hits were only profiled thereafter for activity against additional life cycle stages. Whilst this strategy can identify compounds targeting two or more life cycle stages, it does not allow *de novo* discovery of compounds with selective activity against specific life cycle stages such as gametocytes. Parallel screening against multiple life cycle stages would best identify such compounds. These efforts rely on selective and predictive assays for gametocytocidal activity,^4^ transmission-blocking,^5, 6^ and hepatic development.^7^ Moreover, identifying stage-specific compounds will allow the divergent cell biology associated with the different life cycle states to be targeted.^5, 6, 8^ Recently, parallel screening of diversity sets has resulted in reports of such stage-specific compounds.^7, 9, 10, 11^

The Medicines for Malaria Venture (MMV) Pandemic Response Box (PRB) (in partnership with DNDi) is a collection of 400 drug-like compounds stratified by antibacterial, antiviral or antifungal activity (201, 153 and 46 compounds, respectively), with some compounds having antineoplastic activity. The unique and diverse nature of the compounds in the box allow one to explore and target the unique biology in the different life cycle stages to identify new chemical starting points for antimalarial development. We describe here the parallel screen of the MMV PRB on different life cycle stages of *Plasmodium* including asexual stage parasites, liver stage parasites, mature (stage IV/V) gametocytes, male gametes and oocysts (Figure 1). Finally, active compounds were screened for endectocidal activity against mosquitoes. All screens were performed on the human parasite *P. falciparum*, except for the liver stages, where the established *P. berghei* assay was used.^12, 13^ Hit selection and progression of compounds in our screening cascade was not biased towards activity on any single life cycle stage, allowing the discovery of multistage-active scaffolds and those with stage-specific activity. Importantly, we report the profiling of a subset of compounds as new transmission-blocking molecules that would not have been identified in a test cascade that began solely with an asexual blood stage assay. Four transmission-targeted leads include compounds that are chemically tractable, with good physicochemical properties and novel modes-of-action, amenable to development as transmission-blocking antimalarials.

**Figure 1:**
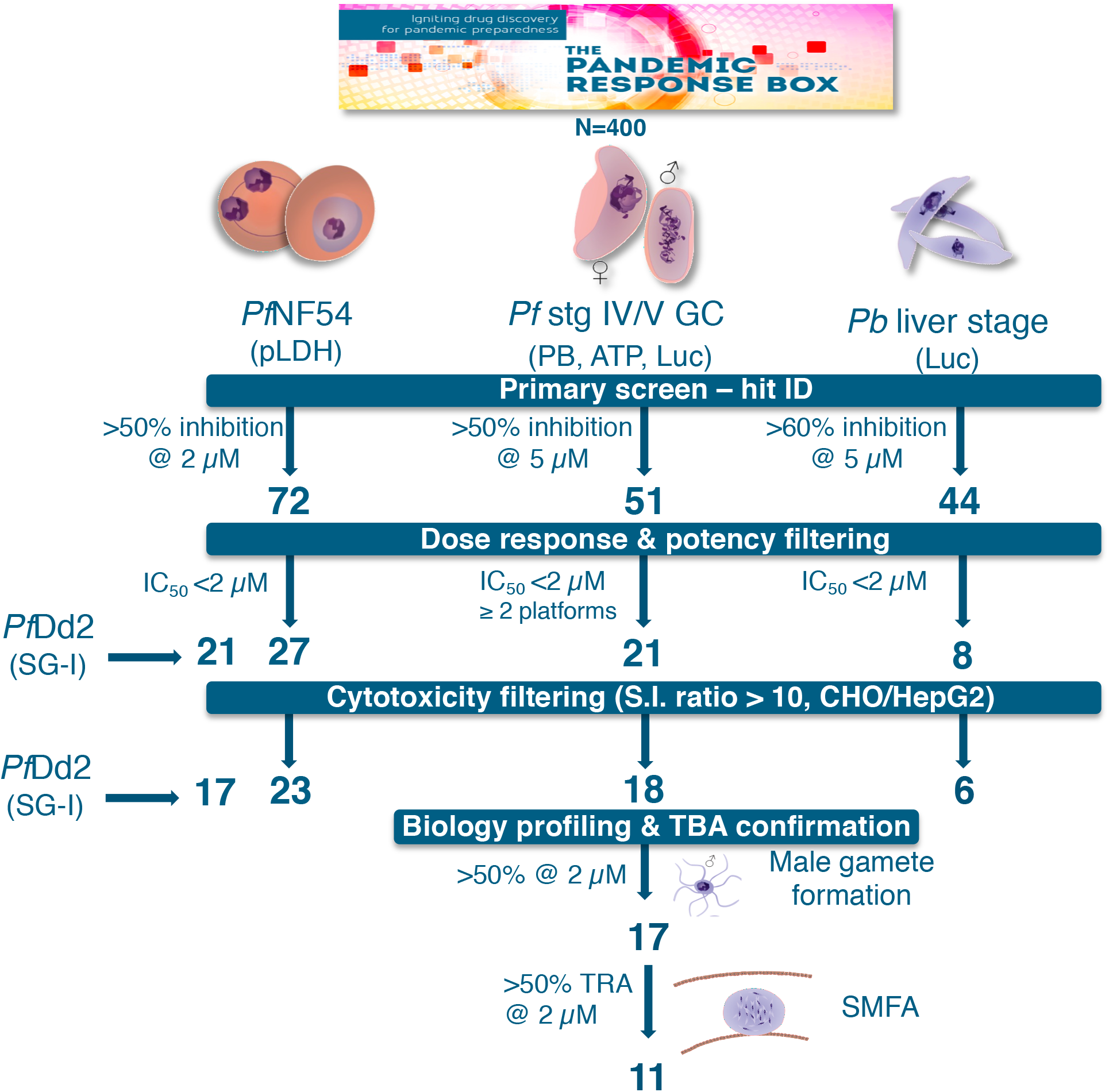
Screening cascade of the MMV Pandemic Response Box for activity against multiple life cycle stages of *Plasmodium*. The 400 compounds in the PRB were screened in a primary assay against drug sensitive (NF54) *P. falciparum* asexual blood stages (ABS, at 2 and 20 μM) and mature gametocytes (stage IV/V, GC, 1 and 5 μM) and *P. berghei* liver stages (5 μM). Hits were selected based on ≥50% inhibition at specific concentrations as indicated. The criteria for each decision point are indicated followed by the number of compounds that passed the criteria. Compounds were additionally evaluated in dose-response on drug resistant asexual Dd2 parasites (chloroquine, pyrimethamine and mefloquine resistant). IC_50_: 50% inhibitory concentration, pLDH: parasite lactate dehydrogenase assay, PB: PrestoBlue^®^ assay, ATP: ATP viability assay, Luc: luciferase reporter lines assays. *Pf: Plasmodium falciparum. Pb: Plasmodium berghei*, S.I.: selectivity index; CHO: Chinese hamster ovarian cells; HepG2: hepatocellular carcinoma line; TBA: transmission-blocking activity; TRA: transmission-reducing activity; SMFA: standard membrane feeding assay.

## RESULTS

### Parallel screening of the PRB reveals hits against multiple life cycle stages

To identify active compounds against different stages of the *P. falciparum* life cycle (irrespective of their activity against the other life cycle stages), the PRB was screened in parallel against asexual *Pf*NF54, stage IV/V gametocytes from *Pf*NF54 and *P. berghei* liver stages (Figure 1). To validate the *Pf*NF54 stage IV/V gametocyte data, we orthogonally screened all compounds on three independent gametocyte assay platforms to confirm that hit selection (compounds active on at least 2 platforms) was independent of assay readout.^4^ Primary hits were identified with a relatively lenient but inclusive cut-off of ≥50% inhibition (at 2 μM for asexual stages and 5 μM for gametocytes and liver stages, the latter at >60% cut-off). Asexual stage activity was confirmed against drug resistant *Pf*Dd2 asexual parasites. Cytotoxicity filtering was applied after evaluation of the IC_50_, and transmission-blocking potential of compounds with gametocytocidal activity was confirmed by inhibition of male gamete exflagellation and in a standard membrane feeding assay (SMFA).

An 18% hit rate was obtained against *Pf*NF54 asexual parasites, 12% against *Pf*NF54 stage IV/V gametocytes and 11% against liver stages (Figure 2A, Supplementary file S1). Although a number of compounds showed activity against all these life cycle stages, stage-specific differentiation was evident, as exemplified by the overrepresentation of antifungal compounds in the hit pool for stage IV/V gametocytes compared to asexual parasites (Figure 2B). The remaining hits reflect the distribution of the compounds in the MMV PRB, with the highest number of hits classified as antibacterials followed by antivirals. The latter seemed to be more potent (as a % of the hits) on *Pf*NF54 stage IV/V gametocytes relative to *Pf*NF54 asexual parasites. Only four compounds showed marked toxicity against CHO cells (<50% viability at 2 μM, supplementary Fig. S1). There is little overlap of the compounds in the box with typical antimalarial scaffolds identified to date, as shown by the superimposition of the PRB chemical space on the current antimalarial drug within the launched drugs chemical space (Supplementary Fig. S2).

**Figure 2:**
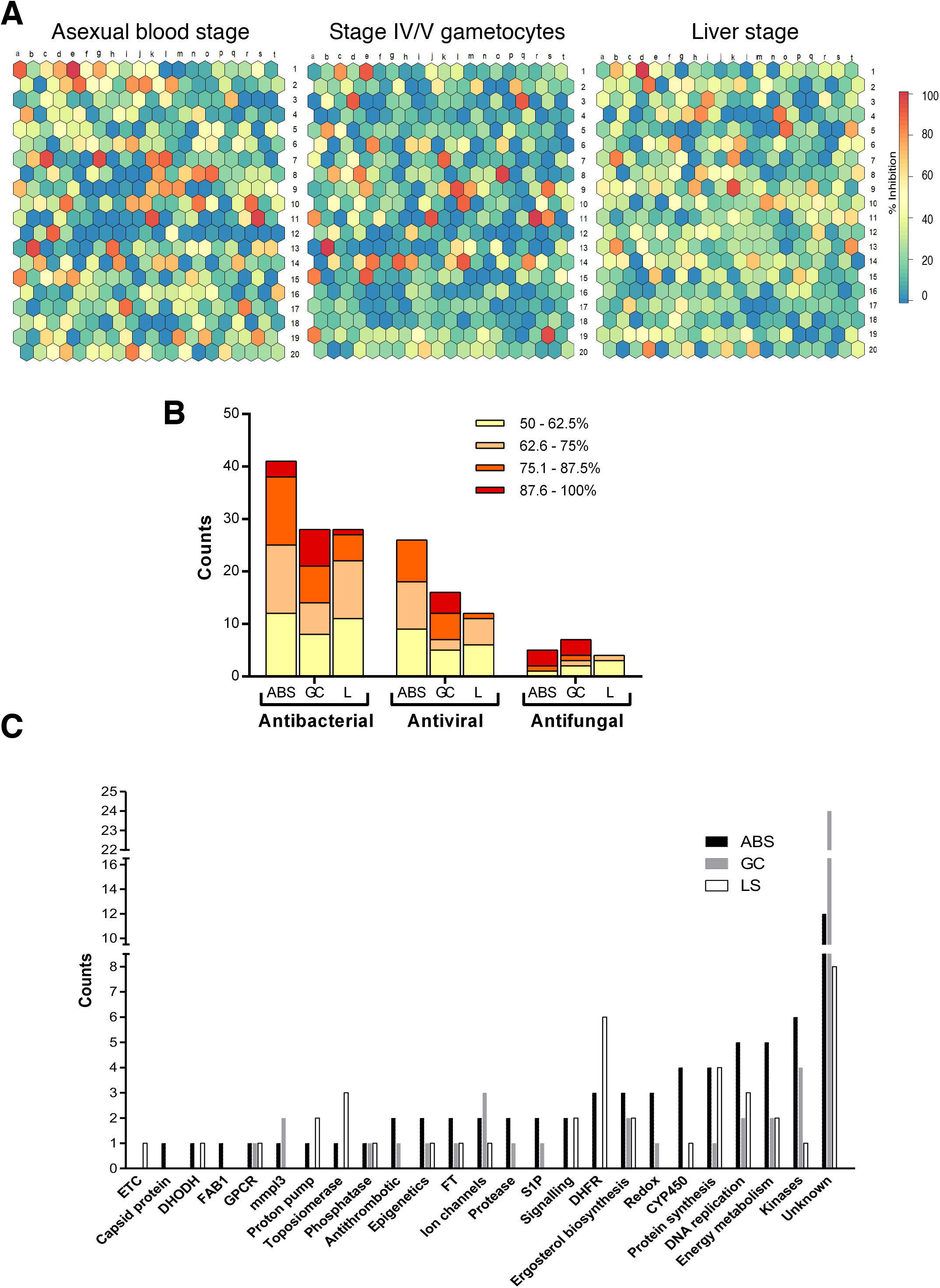
Primary screening of the MMV PRB for hits against *P. falciparum* parasites. **(A)** Supra-hexagonal maps of all 400 compounds included in the MMV PRB after analysis on *P. falciparum* NF54 asexual blood stage parasites and stage IV/V gametocytes. Each hexagon is indicative of a single compound and the order of the hexagon is the same between the two plots. Colors on the heat bar indicate % inhibition of proliferation (asexual blood stage parasites) or viability (stage IV/V gametocytes) after treatment with each compound at either 2 μM (asexual blood stages) or 5 μM (stage IV/V gametocytes or liver stages), screened at least in duplicate. The data for the stage IV/V gametocytes are compiled from hits identified with three different assay platforms, run in parallel (ATP, PrestoBlue^®^ and luciferase reporter expression) with any hit on any platform included, and where identified on >2 platforms, the highest value was included. **(B)** Proportional distribution of hits (>50% inhibition @ 2 μM for asexual blood stages or 5 μM for stage IV/V gametocytes and liver stages) based on disease area as defined in the MMV PRB. Bars are delineated to show activity distribution. ABS = asexual blood stages, GC = stage IV/V gametocytes, L= liver stage. **(C)** Stratification of hits based on biological activity or target indicator. Protein targets / metabolic pathways were identified based on the descriptions of compounds with known activity in the MMV PRB in other disease systems. FT: farnesyltransferase inhibitors; GPCR: G-protein coupled receptors; S1P: sphingosine-1-phosphate receptor modulators, CYP: cytochrome inhibitors; ETC: electron transport chain, DHFR: dihydrofolate reductase, DHODH: dihydroorotate dehydrogenase.

Based on the target indicators / biological pathway descriptors available for the compounds in the PRB in other diseases, *Pf*NF54 asexual hits were enriched for inhibitors of kinases, CYP450, energy metabolism and DNA synthesis. Inhibitors of dihydrofolate reductase (DHFR, antifolates), dihydroorotate dehydrogenase (DHODH), proton pumps and topoisomerase were exclusively hits for *Pf*NF54 asexual parasites and liver stages. Compounds with both *Pf*NF54 asexual and gametocyte activity include antithrombotics, protease inhibitors, sphingosine-1-phosphate receptor modulators and compounds affecting redox homeostasis, whereas inhibitors of MmpL3 (mycobacterial membrane protein large 3) and ion channels were predominant in gametocyte hits (Figure 2C). Interestingly, well known antimicrobials (e.g. thalidomide, isoniazid and saquinavir) were not active in our screens. Chemical classes highly represented in the hit pool include quinolines, benzamides/benzoids and azoles.

### Novel multistage-active compounds

All hit compounds were counter screened against either CHO or HepG2 mammalian cells to remove cytotoxic compounds (supplementary file S2). Of the 72 MMV PRB hits with *Pf*NF54 activity (>50% at 2 μM), 23 were active with IC_50_ values <2 μM. An additional five compounds were active on *Pf*Dd2 (Figure 2A, supplementary file S2). Of these 28 compounds, 16 were exclusively active against the asexual stages (Figure 3A). Of the 51 hits active against *Pf*NF54 stage IV/V gametocytes, 18 compounds had IC_50_ values <2 μM. Eight shared activity against asexual stages but ten had gametocyte stage-specific activity (Figure 3A). Only six compounds showed activity against *P. berghei* liver stages (<2 μM). Notably, two compounds were active (IC_50_ ≤2 μM) against all life cycle stages: the peptidomimetic antitumor agent MMV1557856 (Birinapant), a second mitochondrial-derived activator of caspases (SMAC) mimetic inhibitor of apoptosis protein (IAP) family members^14^; and the imidazoquinoline antitumor agent MMV1580483 (AZD-0156), a DNA-damage signaling kinase inhibitor (Ataxia Telangiectasia Mutated kinase) (Figure 3B).

**Figure 3:**
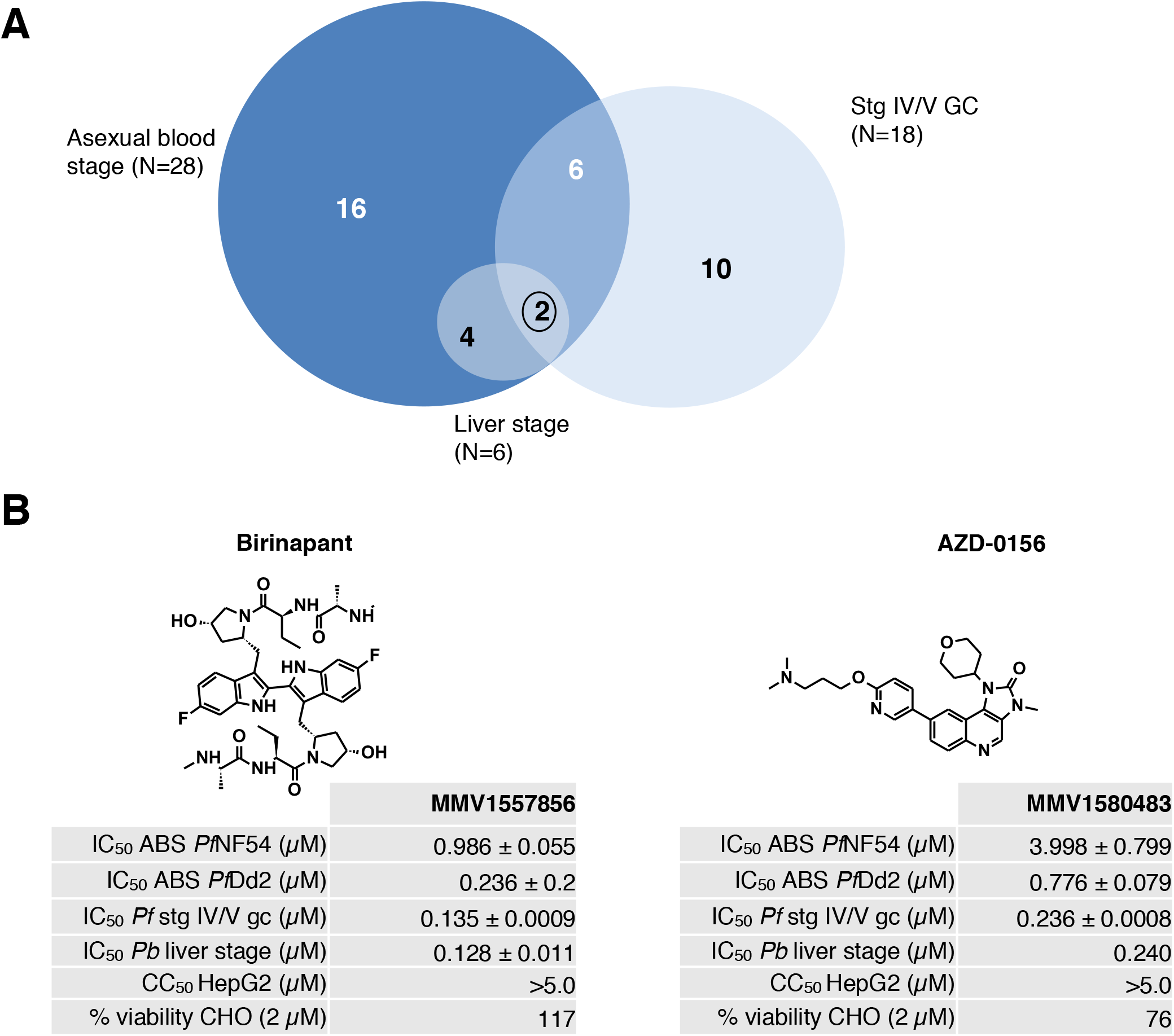
Active compounds on multiple stages of *Plasmodium* development after dose response evaluation and cytotoxicity filtering. **(A)** Venn diagram of the number of compounds identified with activity (inhibitory concentration, IC_50_) below 2 μM, for which no cytotoxicity was identified on either CHO cells (>50% viability at 2 μM or selectivity index >10) or HepG2 cells (selectivity index >10). **(B)** A total of 2 compounds with activity against all life cycle stages tested: Birinapant and AZD-0159. Asexual blood stage activity (ABS) was determined against both drug sensitive (NF54) and drug resistant (Dd2) *P. falciparum*. GC: *P. falciparum* stage IV/V gametocytes. Toxicity indicated both at CC50 (cytotoxic concentration) against HepG2 cells as well as for viability of CHO cells remaining after 2 μM treatment. MMV codes related to compound codes provided in the box. Data are from at least three independent biological repeats, performed with minimum technical duplicates, ± S.E.

### New asexual parasite specific chemotypes

Encouragingly, the 28 compounds with asexual parasite activity (Supplementary file S2) included the known antimalarial compounds chloroquine (MMV000008) and tafenoquine (MMV000043), which were both present in the MMV PRB and showed IC_50_s comparable to those previously reported^6^ (30 nM and 940 nM, respectively), validating the screening process (Figure 4). The most potent of the 28 compounds was an antibacterial diaminopyridine propargyl-linked antifolate,^15^ MMV1580844 (IC_50_ = 0.0017 μM), which targets DHFR in mammalian and yeast cells.^16, 17^ It also showed activity against *P. berghei* liver stages (0.004 μM). As there was no activity against *Pf*NF54 stage IV/V gametocytes, our data concur that inhibition of *Plasmodium* DHFR is only important to asexual and liver stage schizogony.^6^ MMV1580844 had a pronounced (63-fold) loss of activity against the antifolate (pyrimethamine) resistant *Pf*Dd2 line. By contrast, the quinazoline antifolate trimetrexate (MMV1580173, derived from methotrexate) was potently active against *Pf*Dd2 (IC_50_ = 0.108 μM) as well as *P. berghei* liver stages (IC_50_ = 0.0005 μM), in both instances with more than 10-fold selectivity towards the parasite versus CHO cells. As with MMV1580844, MMV1580173 did not display any gametocytocidal activity.

**Figure 4:**
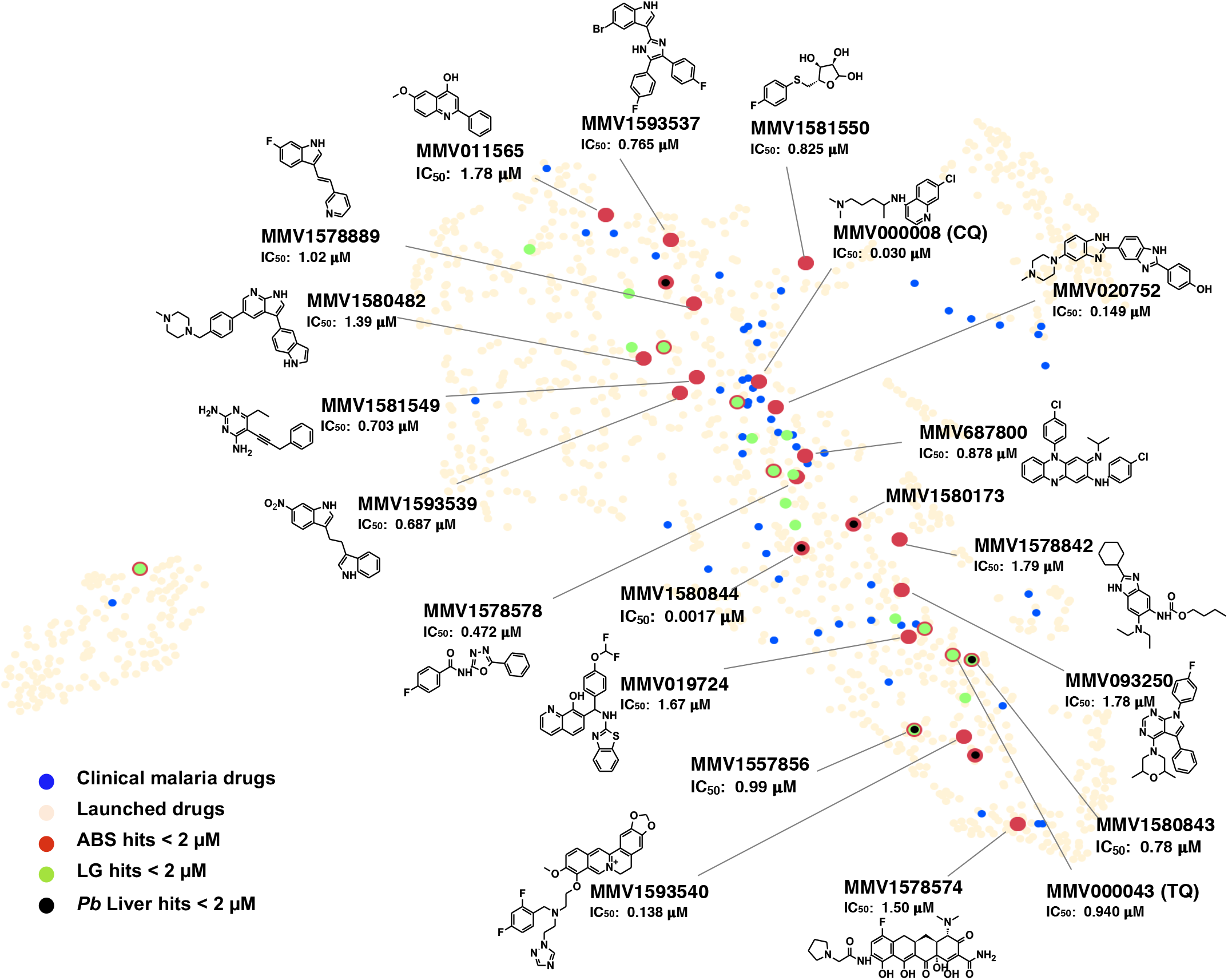
Asexual blood stage active compounds from the MMV PRB in relation to malaria clinical drugs. MMV PRB compounds active against asexual blood stage (ABS) parasites of *P. falciparum*. Compounds with inhibitory concentrations (IC_50_) below 2 μM were identified as hits against either *Pf*NF54 or *Pf*Dd2. MMV PRB hits are represented in the Launched Drugs chemical space (beige) in comparison to Malaria Clinical Drugs (blue dots). Asexual blood stage actives with IC_50_ values <2 μM are indicated in red, gametocyte actives and liver stage actives (all at the same cut-off) are indicated in green and black, respectively, with different dot diameters to highlight compounds active on multiple stages. The 16 asexualspecific compounds are labelled with Compound ID, asexual stage IC_50_ and structure and other compounds of interest just by name and IC_50_. CQ: chloroquine, TQ: tafenoquine.

The majority (13/16) of the asexual-specific compounds have not been previously reported with antiplasmodial activity, nor do they show structural similarity to any other compounds with antiplasmodial activity, as highlighted by the absence of overlap between the two chemical spaces (hits and antimalarial drugs) (Figure 4, supplementary Fig. S2). Activity was confirmed for MMV687800 (Clofazimine, IC_50_ = 0.88 μM), MMV1578574 (eravacycline, IC_50_ = 1.5 μM) and MMV011565 (IC_50_ = 1.78 μM), all with previously described antiplasmodial activity in PubChem (https://pubchem.ncbi.nlm.nih.gov/). Interestingly, the majority of the asexual-specific compounds are classified as antibacterials and include two kinase inhibitors, MMV1593539 (IC_50_ = 0.686 μM), a pyruvate kinase inhibitor, and MMV1580482 (URMC-099-C, IC_50_ = 1.3 μM), a mixed lineage kinase 3 (MLK3) inhibitor. Additionally, there is pibenzimol (MMV020752, IC_50_ = 0.149 μM) a disrupter of DNA replication and MMV019724 (IC_50_ = 1.67 μM), an antiviral lactate dehydrogenase inhibitor.

### Liver stage activity is associated with asexual parasite activity

Six dual stage compounds (asexual blood stage and *P. berghei* liver stage activity ≤ 2 μM) were identified in the MMV PRB, marking them as having prophylactic potential (IC_50_ range from 0.0005 – 1.72 μM, Figure 4). These compounds include previously described antifolates (MMV1580173 and MMV1580844) but also the ribonucleotide reductase inhibitor MMV1580496 (triapine) and the bacterial methionyl-tRNA synthetase inhibitor MMV1578884 (REP3123) in addition to the multistage-active compounds, AZD-0156 and Birinapant.

### Unique compounds with stage-specific activity against IV/V gametocytes

Eighteen compounds were active (IC_50_s <2 μM) against late stage gametocytes, without showing toxicity to mammalian cells, as confirmed in at least two of the three orthogonal assays (ATP, PrestoBlue^®^ or luciferase reporter assays, Supplementary file S2). Of these, eight compounds shared asexual parasite activity, but more importantly, ten compounds selectively inhibited *Pf*NF54 stage IV/V gametocyte viability (>10-fold difference in IC_50_ between gametocytocidal and asexual activity) with IC_50_s <0.5 μM (Figure 5). The most potent compound was the antineoplastic epidrug ML324^18^ (MMV1580488, IC_50_ = 0.077 μM) that targets jumonji domain demethylases (KDM4). There was also a marked selection for structurally unrelated compounds that bind G-protein coupled receptors (GPCRs) and related transmembrane proteins (Figure 5A). These include MMV1581558 (IC_50_ = 0.130 μM) and two inhibitors of MmpL3: the well characterized 1,2-ethylene diamine antitubercular clinical candidate SQ109 ^19, 20, 21^ (MMV687273, IC_50_ = 0.105 μM) and a rimonabant derivative MMV1580843^22^ (IC_50_ = 0.108 μM). Since MmpL3 inhibitors act rapidly in mycobacteria and trypanosomatids,^21, 23^ we evaluated the gametocytocidal action of MMV687273 and MMV1580843 for 12, 24 and 48 h (Figure 5B), with no significant difference (*p*=0.937 and *p*=0.558, respectively; one-way ANOVA, DF=2, n=3) observed in the IC_50_ values, indicating activity within 12 h of exposure to gametocytes. Moreover, the activity of MMV687273 (SQ109) was similar to *S,S*-ethambutol.2HCl^20^ (MMV687801) whereas 2-adamantanamine (MMV180402) was inactive (Figure 5C).

**Figure 5:**
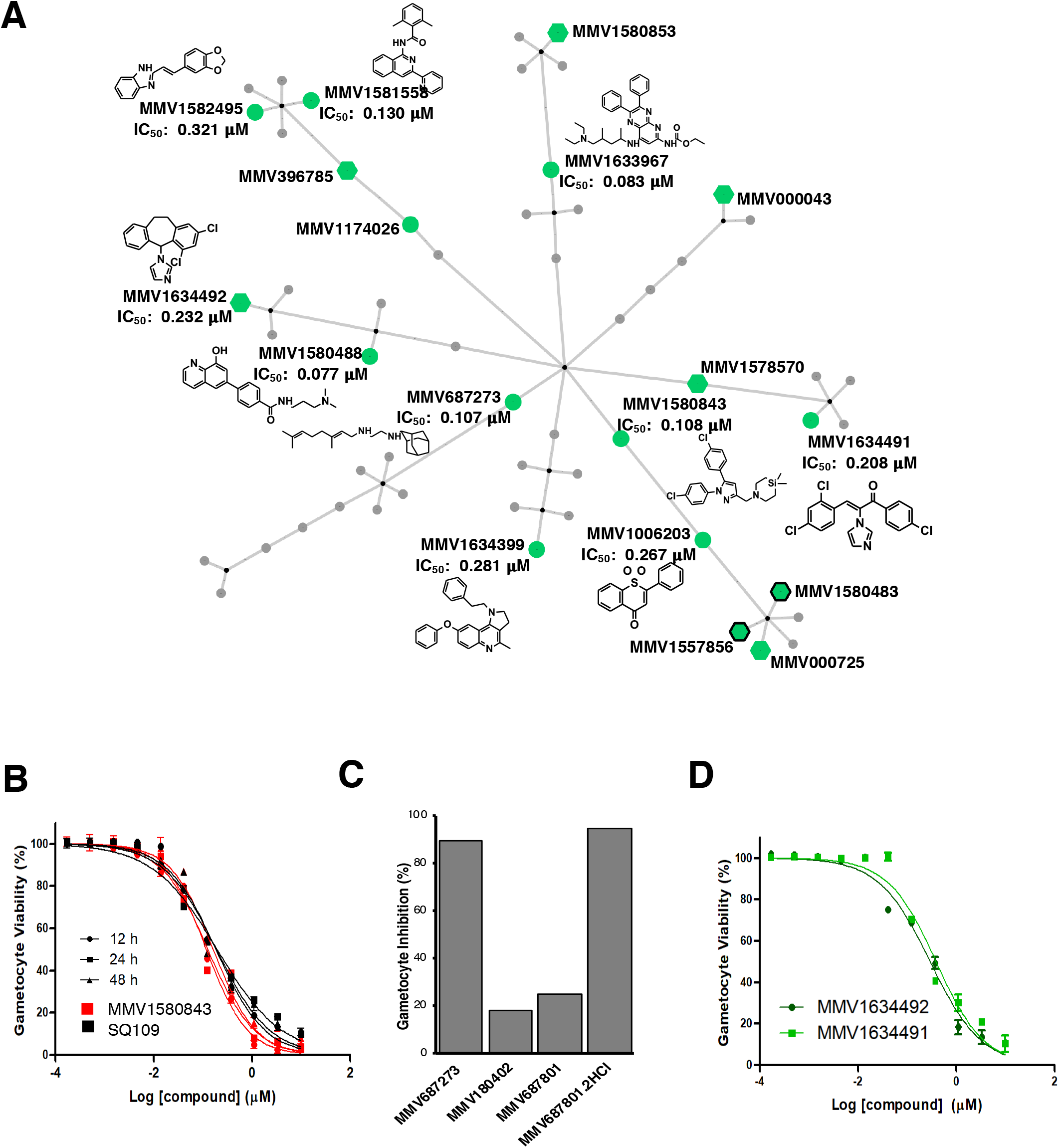
Active compounds from the MMV PRB against *P. falciparum* stage IV/V gametocytes. **(A)** Chemical cluster analysis of the gametocyte hit compounds, using the *FragFP* descriptor and a Tanimoto similarity index >0.50 in OSIRIS DataWarrior v 5.0.0, and network construction with Cytoscape v 3.7.2. Edges were assigned between similar scaffolds and a parent node. Active compounds with IC_50_ values <2 μM are indicated in green, those with additional activity at the same cut-off on ABS are indicated with hexagons and those with shared activity on liver stages with black borders. Structures are highlighted for selected compounds. **(B)** Dose-response of MmpL3 type inhibitors MMV687273 (SQ109, black lines) and the rimonabant derivative MMV1580843 (grey lines), tested on stage IV/V gametocytes after 12, 24 or 48 h drug pressure (PrestoBlue^®^ assay). **(C)** The % inhibition of stage IV/V viability under 2 μM pressure of MMV687273 (SQ109) compared to MMV180402 (2-adamantanamine) or the parent compound ethambutol (MMV687801, *S,S*-ethambutol and its HCl derivative). **(D)** Gametocyte-specific compounds were enriched for azole antifungals, including MMV1634491 and MMV1634492, with dose-responses of these two compounds indicated after 48 h exposure to stage IV/V gametocytes and evaluated on the PrestoBlue^®^ assay. Data are from three independent biological repeats, each performed in technical triplicates, ± S.E.

Two imidazole antifungals showed potent activity against gametocytes (Figure 5D), though they differed structurally. MMV1634491 was >10-fold more active against gametocytes (IC_50_ = 0.208 μM) than against *Pf*NF54 asexual parasites (IC_50_ = 2.6 μM) while MMV1634492, the topical antifungal eberconazole, showed more equipotent activity (IC_50_s = 0.23 and 0.15 μM, respectively). Eberconazole targets sterol biosynthesis by inhibiting lanosterol 14α-demethylase CYP51^24^, as likely does MMV1634491 based on structural similarity with other azole antifungals.^25^

### Gametocytocidal compounds target male gametes

The transmission-blocking activity of gametocytocidal hits was validated on male gametes, as these display increased sensitivity to compounds.^26^ A functional male gamete exflagellation inhibition assay was performed in carry-over format, where compounds (2 μM) were added to mature gametocytes for 48 h before male gamete exflagellation was induced and measured ^27^ (Figure 6A). The majority of compounds (14) inhibited male gamete exflagellation by >60%, 11 of which were potent at ≥80% inhibition. The latter included the gametocyte-targeted compounds MMV1580488 (ML324), the azole antifungals MMV1634491 and MMV1634492 and the MmpL3 inhibitors MMV1580843 and MMV687273, as well as compounds with additional asexual parasite activity, i.e. MMV1580483, MMV396785, MMV1582495, MMV1578570 and MMV1581558.

**Figure 6:**
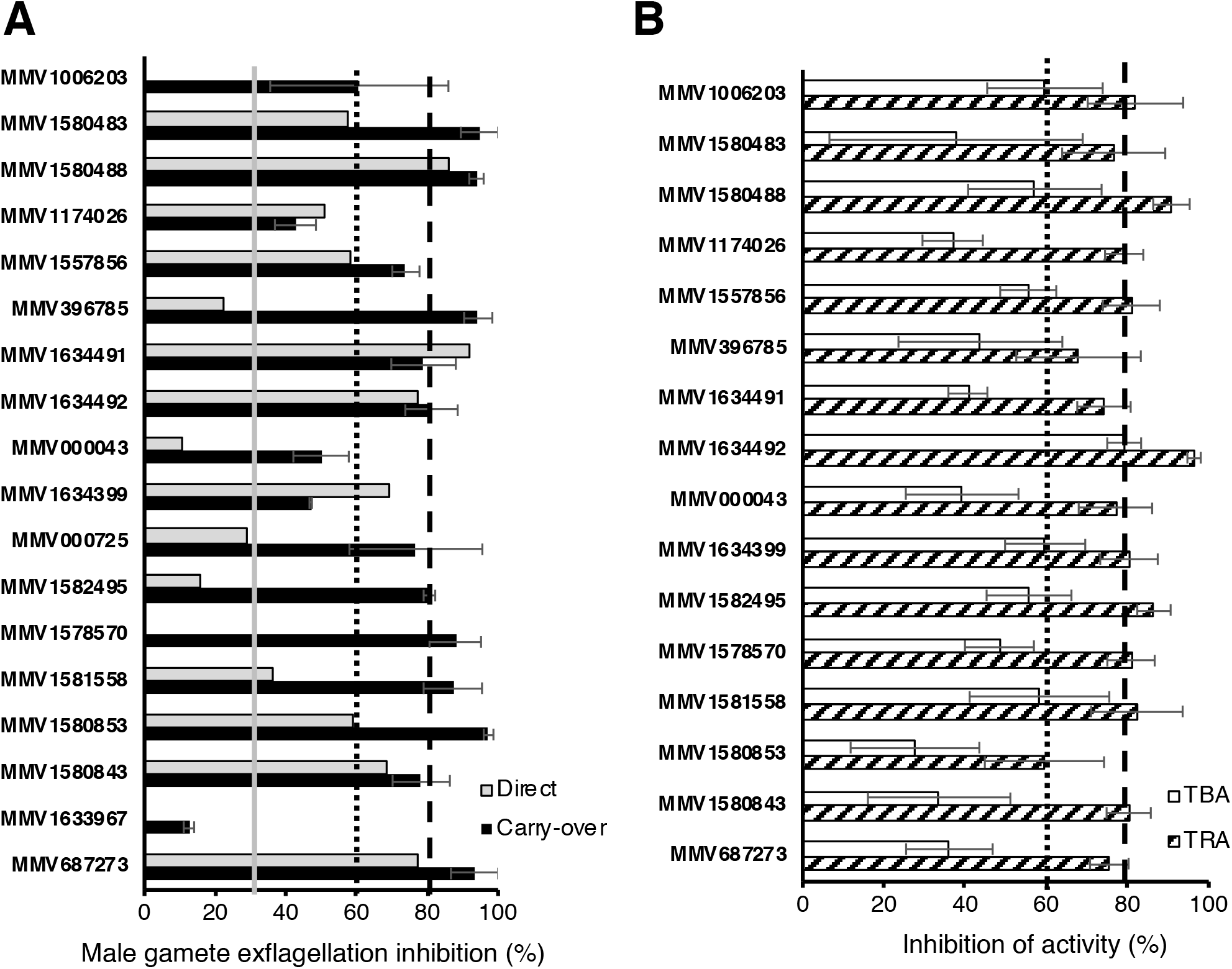
Confirming the transmission-blocking activity of gametocytocidal compounds in the MMV PRB. **(A)** 18 compounds with activity against *P. falciparum* stage IV/V gametocytes were evaluated for their ability to inhibit male gamete exflagellation. Compounds (2 μM) were either used on stage IV/V gametocytes for a 48 h treatment prior to inducing male gamete exflagellation (carry-over format) or were directly added during induction of exflagellation (20 min incubation). Exflagellation was induced with 100 μM xanthurenic acid and 20% human serum, (A+ male) and exflagellating centers semi-automatically quantified from 15 videos of 8-10 seconds each 15 and 22.5 min after incubation, under 10x magnification. Data are from at least two independent biological repeats, performed in technical triplicates, ± S.E. **(B)** SMFA data for 16 compounds (selected based on >50% inhibition on male gamete exflagellation). SMFA was performed by feeding *An. coluzzii* mosquitoes with compound-treated gametocyte cultures (48 h treatment at 2 μM). Data are presented as % of TRA (transmission-reducing activity, reduction in oocyst intensity) or % TBA (transmission-blocking activity, reduction in prevalence) from at least three independent biological repeats, performed with technical duplicates, ± S.E.

To determine if the compounds also directly targeted gametes, the assay was repeated both in ‘washout’ format (48 h drug pressure on gametocytes, after which male gamete exflagellation was induced in the absence of compound) and in a ‘direct’ format, where compounds were only added during exflagellation induction.^27^ All the compounds except for MMV1578570 retained activity in the washout experiment, indicating that their gametocytocidal activity was irreversible (Supplementary Fig. S3). Five compounds (the epidrug MMV1580488, the two azole antifungals MMV1634491, MMV1634492, a quinoline MMV1634399, and the two MmpL3 inhibitors MMV1580843, MMV687273) retained activity on male gametes in the direct format, implying that shared essential biology between these stages is being targeted (Figure 6A).

### Novel chemotypes with confirmed transmission-blocking activity

The final confirmatory step of transmission-blocking is the mosquito-based SMFA,^28, 29, 30^ which was performed using an African malaria vector, *An. coluzzii* (G3). Females were fed on a blood meal infected with *Pf*NF54 late stage gametocytes and treated with selected compounds that displayed >50% male gamete exflagellation activity at 2 μM. TRA (reduction in oocyst intensity) and TBA (reduction in oocyst prevalence) were determined after 8-10 days (Figure 6B, supplementary File S3). Total sample size for the control feeds averaged at 53 mosquitoes, with average oocyst prevalence at 71% and oocyst intensity of 5.8 oocysts/midgut. The TRA for MMV000043 (Tafenoquine) at 77% correlates well with previous reports on *An. stephensi*,^31^ validating assay performance with *An. coluzzii*. All the compounds evaluated reduced TRA by >60% (except for MMV1580853) and, remarkably, nine compounds had ≥80% TRA. Four gametocyte-targeted compounds MMV1006203 (1,1-dioxide 1-Thioflavone), the azole antifungal MMV1634492, a quinoline MMV1634399 and the GPCR inhibitor MMV1581558 were able to block transmission (TBA) by ≥60% (MMV1634492 by 79%), associated with a significant reduction in oocyst intensity (*p*<0.05, supplementary File S3) of these structurally dissimilar compounds.

### Endectocidal activity

The compounds with transmission-blocking potential were additionally evaluated for their activity as endectocides, killing mosquitoes after being supplied in a blood meal. However, none of the compounds produced significant mortality (one-way ANOVA, *p* = 0.7005, total DF=71, n≥2) in the 4-day mortality assay at 2 μM compared to DMSO as control under these conditions (supplementary Fig. S4). Rather, moderate killing (~30%) was observed for the two MmpL3 inhibitors MMV687273 (SQ109) and MMV1580843, the GPCR inhibitor MMV1581558, AZD-0156 (MMV1580483) and MMV1174026, marking these compounds with a potential to kill the parasite as well as the mosquito vector. Interestingly, all these compounds also had >60% reduction of TRA in addition to their potential endectocidal activity.

## DISCUSSION

The ability to quickly respond to pandemics has become of paramount importance, and compound sets like the MMV PRB provide an essential tool to support rapid screening of diverse druggable compounds for potential repurposing. Indeed, antimalarials have previously been investigated as antineoplastics^32, 33^ and hydroxychloroquine/chloroquine are currently undergoing clinical evaluation for the treatment of Covid-19. Conversely, several antibiotics and antifungals have previously demonstrated antimalarial activities.^34, 35^ Here, we screened the MMV PRB across multiple *Plasmodium* stages and identified chemical matter with antimalarial activity not previously described, providing a useful resource to the research community for drug repurposing.

Multistage activity is a preferential attribute for the next generation of antimalarials,^36^ but such compounds are rarely found in diversity library screens, in large part due to targeted screening approaches rather than parallel screening in multiple assays. We identified two non-cytotoxic multistage-active compounds in the PRB (Birinapant and AZD-0156), that could point to biological parsimony of conserved targets in all these stages, essential to the survival of the parasite. Both compounds inhibit proteins involved in cellular stress responses by either inducing apoptosis or preventing DNA damage recovery responses. Interestingly, Birinapant has recently been shown to preferentially kill *Plasmodium-infected* hepatocytes (attributed to reducing host cellular IAP) but did not affect asexual stage parasitemia in a *P. berghei* mouse model.^37^ It would be imperative to determine what proteins are targeted by these compounds in the different life cycle stages and to understand the reasons for the differences in parasitemia seen *in vitro* herein and *in vivo* previously.

The dual-active asexual and liver stage compounds identified in the PRB have the potential for prophylactic and chemoprotective utility (target candidate profile 4, TCP-4), in addition to being chemotherapeutically relevant (target candidate profile 1, TCP-1).^36^ Though compounds targeting the same parasite protein in both liver and asexual stages have the associated risk of target-based resistance, the smaller number of parasites in the liver stage reduces this risk. However, with compounds like the ribonucleotide reductase inhibitor MMV1580496 (triapine), there is the added issue of cytotoxicity risks since, as anticancer agent, it invokes DNA biosynthesis as a pathway. Thus, it is important to identify the mechanism-of-action of these compounds with respect to the liver stage of the parasite. For instance, the dual-active MMV1578884 (REP3123) is a *Clostridium difficile* methionyl-tRNA synthetase (metRS) inhibitor and could target aminoacyl-tRNA synthetases (aaRS) in *P. falciparum*. To our knowledge there is no aaRS inhibitor in clinical development as antimalarial but structural differences between several *Pf*aaRS and their human counterparts are encouraging that selectivity can be achieved.^38^ Notably, MMV1578574 (eravacycline), as a tetracycline-class antibacterial recently approved for the treatment of complicated intra-abdominal bacterial infections, binds to the 30S ribosomal subunit similar doxycycline, the latter used prophylactically for malaria and demonstrating liver stage activity clinically.^34^ Eravacycline was only identified with asexual stage activity in our screens and not liver stage activity.

The involvement of protein and lipid kinases in key pathogen functions have made inhibitors thereof a focus of drug design strategies including those that affect multiple life cycle stages.^39^ Amongst the asexual stage active compounds identified, MMV1580482 (URMC-099) operates as a human MLK3 inhibitor and MMV1593539 as a *Staphylococcus aureus* pyruvate kinase inhibitor. However, due to MMV1580482’s activity against breast cancer cell lines and brain metastatic variants while notably having a brain-penetrant capability, achieving *Plasmodium* selectivity in an analogue program directed against malaria will be challenging. In addition to selectivity over human homologues, MMV1593539 also contains a nitro group that is considered a red flag, and safety needs to be assessed or attempts made to replace this group. Interestingly, of the two reported pyruvate kinases in *P. falciparum* (PK1 and PK2), PK1 has a crystal structure (<u>10.2210/pdb3KHD/pdb</u>), which could guide selectivity and optimization studies provided *Pf*PK1 is responsible for phenotypic activity.

Importantly, our parallel screening approach on different life cycle stages yielded compounds and chemical scaffolds that not only have stage-specific asexual parasite activity but also selectively and specifically target the elusive gametocyte stages with activity in mosquito transmission assays. This unbiased approach, where compounds are not only profiled for additional life cycle activity once asexual activity has been established, confirms the possibility of identifying gametocyte-specific compounds.^7, 9, 11^ Indeed, we identified several active compounds that have no previous documentation of antimalarial activity, simply because they were not screened against the correct life cycle form of the parasite where the relevant biology being targeted was essential.

Our stringent profiling cascade additionally ensured a high success rate in confirming transmission-blocking activity and validates the use of orthogonal gametocytocidal screens^4, 11^ as a primary filter in large scale screens. In addition, this approach resulted in a linear correlation between gametocytocidal activity and activity against male gametes, which directly translated to oocyst reduction. By evaluating both TRA and TBA, our data highlighted the importance of both parameters in evaluating SMFA data. Importantly, we showed a large reduction in TRA for some compounds. This indicates that the decreased number of oocysts carried by such mosquitoes will result in a lowered efficacy of transmission, as intensity of a mosquito infection has been shown to be critically important to the success of transmission.^40^ However, we additionally observe a decrease in oocyst prevalence (TBA), that implies that the majority of mosquitoes treated with compounds that affect oocyst prevalence would not carry parasites. The latter will also therefore have an epidemiological impact in line with WHO-recommended vector control interventions, where an efficacy of >80% would result in a major impact on transmission. Interestingly, three of the most active transmission-blocking compounds with epidemiological impact also somewhat affect the mosquito vector themselves. These compounds could indeed provide a conceptual innovation, as they would fit both TCP-5 and −6 criteria and therefore block transmission as well as shorten the lifespan of the mosquito,^41^ particularly if their potency against the mosquito could be improved.

Gametocyte-targeted compounds have become important to deliver leads for development as transmission-blocking specific antimalarials, filling the niche required for TCP-5.^36^ Compounds with selective transmission-blocking activity would presumably target divergent biological processes compared to those in asexual parasites^42^ and this, in addition to the low parasite numbers in transmission stages and non-proliferative nature of gametocytes, would reduce the risk of resistance development. When used in combination with a TCP-1 candidate, such TCP-5 targeted compounds could also protect the TCP-1 drug from resistance development. Alternatively, when used as a stand-alone drug, TCP-5 targeted compounds would decrease the gametocyte burden in the human population, which would be particularly important in pre-elimination settings as add-ons to enhance standard measures of malaria control.

Our data also indicate specific gametocyte-associated biological processes worthy of further investigation. Though the PRB contained nine azole antifungals (including miconazole, ketoconazole, fluconazole), only two imidazoles (eberconazole, MMV1634492, and MMV1634491) showed activity against gametocytes, the latter with >10-fold selectivity relative to the asexual stage. This is the first report of the antimalarial activity of eberconazole, though this has been seen for other azole antifungals. Ketoconazole, miconazole and clotrimazole, like eberconazole, inhibit fungal ergosterol biosynthesis,^24^ but they are also known to target the heme detoxification system in asexual *P. falciparum.^43^* Although heme catabolism is essentially absent in mature gametocytes,^5^ some heme metabolic activity seems to be present in *P. berghei* mosquito stages^44^ and this process as a target for these compounds cannot be excluded. Moreover, as these compounds have exclusive transmission-blocking activity, a novel mode of action might be indicated.

In addition to the azoles, we identified ML324 with exclusive activity against gametocytes. This JmjC demethylase inhibitor was recently shown to be more active against immature gametocytes (~1 μM) than asexual parasites.^45^ Our data showed for the first time that ML324 has increased potency as gametocytes mature to stage IV/V (0.077 μM) and potently kills male gametes with confirmed transmission-blocking activity. This implies that mature gametocytes and male gametes are even more sensitive to changes in histone methylation status due to ML324 treatment, which results in deregulated gene expression, similar to what is seen for other JmjC demethylase inhibitors.^45^

Lastly, the selective transmission-blocking actives included two compounds that are established inhibitors of MmpL3 in bacteria.^19, 20, 21^ Albeit structurally dissimilar, both compounds inhibit MmpL3 through interaction with the protein pore section as indicated by co-crystallization data.^46^ A homologue for this protein is not detectable in the *Plasmodium* genome but the potent (0.107 μM) and selective mechanism-of-action on the non-proliferative differentiated *P. falciparum* gametocytes may be similar to that observed in *Trypanosoma cruzi*, where activity was seen against the non-proliferative and transmissible trypomastigotes (0.05 μM). With the absence of MmpL3 homologues in *T. cruzi*, the activity was explained to be due to the disruption of the proton gradient across the parasite’s mitochondrial membrane.^23^ The possibility of a similar action against *P. falciparum* gametocytes is currently being investigated, in light of the known increased reliance on mitochondrial metabolism compared to asexual *P. falciparum* parasites.^42, 47^ However, the possibility also exists that these compounds interfere with lipid metabolism/transport, which is essential to gametocytogenesis^48^ and oocyst development.^49^ SQ109 and another GPCR inhibitor (MMV1581558) also have the potential to block *P. falciparum* transmission whilst simultaneously affecting the mosquito vector.

An open challenge remains the determination of the mechanism-of-action of transmission-blocking targeted compounds. Since established forward genetic mutant generation routes to identify drug targets are not applicable to the non-proliferating gametocytes, alternative proteomic techniques such as thermal shift, protease protection assays, chemical pull-downs or other technologies available through consortia like MalDA (Malaria Drug Accelerator) will be critical to progress these compounds from discovery to development. One advantage of screening boxes such as the PRB that contain a large number of compounds with already well described DMPK profiles and empirically determined physical characteristics, is that they could progress rapidly through the drug discovery pipeline.

## METHODS

### Ethics statement

This work holds ethical approval from the University of Pretoria Health Sciences Ethics Committee (506/2018); University of Cape Town: AEC017/026; University of the Witwatersrand Human Research Ethics Committee (M130569) and Animal Ethics Committee (20190701-7O); CSIR Research Ethics Committee (Ref 10/2011) and Scripps Research’s Normal Blood Donor Service (NBDS), with approval under IRB Number 125933.

### Parasite culturing

*P. falciparum* asexual parasite cultures, drug sensitive strain NF54 (*Pf*NF54), drug resistant strain Dd2 (*Pf*Dd2, chloroquine, pyrimethamine and mefloquine resistant) and the luciferase reporter line NF54-*Pfs16*-GFP-Luc (kind gift from David Fidock, Columbia University, USA)^50^ were maintained at 37°C in human erythrocytes (5% hematocrit) in complete culture medium RPMI 1640 medium [25 mM HEPES, 0.2% (w/v) D-glucose, ~200 μM hypoxanthine, 0.2% (w/v) sodium bicarbonate, 24 μg/mL gentamicin and 0.5% (w/v) AlbuMAX II lipid-rich BSA or 4.3% (v/v) heat inactivated O^+^ human serum] under hypoxic conditions as previously described.^4^ Gametocytogenesis was induced from asexual parasites (0.5% parasitemia, 6% hematocrit) NF54-background parasites as described.^4^ After a drop in hematocrit (to 4%) after three days, gametocytogenesis was monitored microscopically with daily medium changes. On days 5-10, residual asexual parasites were eliminated with 50 mM N-acetyl glucosamine (NAG) treatment in complete culture medium.

### Asexual blood stage screening

*Pf*NF54 asexual parasite activity was determined with the parasite lactate dehydrogenase assay (pLDH) as described.^51, 52^ All compounds were screened (2 μM and 20 μM) in two replicates on ring-stage cultures (1% hematocrit, 2% parasitemia) for 72 h under hypoxic conditions at 37°C, and survival determined colorimetrically at 620 nm. The IC_50_ (half-maximal concentration) was determined for active compounds under the same conditions. Chloroquine and artesunate were used as controls.

*Pf*Dd2 asexual parasites activity was determined with SYBR Green I as described^13^ on parasite suspensions (0.3% parasitemia, 2.5% hematocrit) in black, clear bottom plates with pre-spotted compounds and incubated at 37°C for 72 h under hypoxic conditions. 2 μL 10x SYBR Green I (Invitrogen) in Lysis buffer (20mM Tris/HCl, 5mM EDTA, 0.16% (w/v) Saponin, 1.6% (v/v) Triton X) was added and plates incubated in the dark at room temperature (RT) for 24 h. Fluorescence was measured using the EnVision^®^ Multilabel Reader (PerkinElmer) (485 nm excitation, 530 nm emission). IC_50_ values were determined in CDD Vault (https://www.collaborativedrug.com/) normalized to maximum and minimum inhibition levels for the positive (Artemisinin) and negative (DMSO) control wells.

### Gametocyte screening^4^

#### PrestoBlue^®^ fluorescence assay

Stage IV/V gametocyte cultures (2% gametocytemia, 5% hematocrit, 100 μL/well) were exposed to compounds and incubated at 37°C for 48 h under hypoxic conditions, stationary, after which 10 μL of PrestoBlue^®^ reagent was added to each well and incubated at 37°C for 2 h. Fluorescence was detected in the supernatant (70 μL, 535 nm excitation, 612 nm emission) with a Tecan Infinite F500 Multimode reader. Dihydroartemisinin (DHA) was used as positive kill control.

#### ATP bioluminescence assay

Stage IV/V gametocyte cultures were enriched with density gradients as described ^4^ and 75 000 gametocytes were seeded into 96-well plates in the presence of compound and incubated for 24 h at 37°C. ATP levels were determined as described with a Promega BacTiter-Glo™ Bioluminescence system.^4^ Methylene blue (MB) was used as positive kill control.

#### Luciferase reporter assay

Stage IV/V gametocytes (2% gametocytemia, 2% hematocrit) from the NF54-*pfs16*-GFP-Luc line ^50^ were seeded with compounds and incubated for 48 h under hypoxic conditions at 37°C. Luciferase activity was determined in 30 μL parasite lysates by adding 30 μL luciferin substrate (Promega Luciferase Assay System) bioluminescence detected with a 10 s integration constant (GloMax^®^-Multi+ Detection System).^4^ MB and MMV390048^53^ were used as positive controls and IC_50_ determined with non-linear curve fitting (GraphPad Prism 6) normalized to maximum and minimum inhibition (DMSO control wells).

### Male gamete exflagellation inhibition assay (EIA)^54^

Gametogenesis was induced on >98% stage V gametocyte cultures by treating with 100 μM xanthurenic acid (XA) in ookinete medium (RPMI 1640 with 25 mM HEPES, 0.2% sodium bicarbonate, pH 8.0, and 20% human serum, A+ male) followed by a >15 min incubation at RT. The EIA was performed as described^54^ on >95% stage V gametocytes, resuspended in 30 μL ookinete medium (culture medium with 100 μM XA, 20% human serum, A+ male). Exflagellating centers were recorded by video microscopy (Carl Zeiss NT 6V/10W Stab microscope with a MicroCapture camera, 10X magnification) in 10 μL activated culture settled in a Neubauer chamber at RT. Centers were semi-automatically quantified from 15 randomly located videos of 8-10 s each after 15-22.5 min.

### *P. berghei* liver stage assay

Potential causal prophylactic activity was tested as previously described ^13^ on HepG2-A16-CD81 cells exposed to compounds and incubated for 24 h. Thereafter, fresh *P. berghei* sporozoites (*P. berghei* ANKA GFP-Luc-SMcon) from infected *An. stephensi* mosquitoes’ salivary gland dissections were added (1 x 10^3^/well), centrifuged (5 min, 330*g*) and incubated at 37°C for 48 h. 2 μL of luciferin reagent (Promega BrightGlo) was added and luciferase activity detected (Perkin Elmer Envision). IC_50_ values were determined in CDD Vault as above (GNF179 as positive, DMSO as negative controls).

### Cytotoxicity counter-screening

Cytotoxicity against Chinese hamster ovarian (CHO) cells was colorimetrically determined using 3-(4,5-dimethyl-thiazol-2-yl)-2,5-diphenyltetrazo-lium bromide (MTT).^55^ Cells were seeded at a density of 106 cells/well in 200 μL medium, incubated 24 h and medium replaced with 200 μL of fresh medium containing the compounds (2 or 20 μM) and incubated for 48 h. Emetine was used as the control. Cytotoxicity against a Hepatocellular carcinoma line (HepG2-A16-CD81) was performed by adding Promega CellTiter-Glo^®^ (2 μL) to these cells seeded as above for the sporozoite liver stage assay but in the absence of sporozoites, and luminescence measured. Curve fitting was done as above using puromycin (positive control) and DMSO (negative control).

### Mosquito rearing and standard membrane feeding assay (SMFA)

*Anopheles coluzzii* mosquitoes (G3 colony, species confirmed in^56, 57^) were reared at 80% humidity, 25°C, 12 h day/night cycle with 45 min dusk/dawn transitions^58^ on a 10% sucrose solution diet supplemented with 0.05% 4-aminobenzoic acid. A mature stage V gametocyte (NF54-strain, 1.5-2.5% gametocytemia, 50% hematocrit, A+ male serum with fresh erythrocytes) was evaluated for male gamete exflagellation and male:female ratio of 1:3 confirmed before proceeding with feeding. Gametocytes were treated with 2 μM of each compound for 48 h. Glass feeders covered with cow intestine with 1 mL of the gametocyte culture, on top of feeding cups (350 mL), were used for SMFA. Each cup contained 25 unfed (2-3 h starvation) *An. coluzzii* females (5-7 days old), fed in the dark for 40 min at RT. After removing unfed/partially fed mosquitoes, females were housed as above for 8-10 days and then dissected to remove midguts, which were rinsed in PBS, incubated in 0.1% (v/v) mercurochrome for 8-10 min and oocysts counted under bright field illumination (20-40x magnification). Reduction in prevalence (transmission-blocking activity, %TBA: 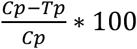, where *p*: oocyst prevalence, *C*: control and *T*: treated) and reduction in number of oocysts intensity (transmission-reducing activity, %TRA: 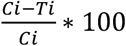, where *i:* oocyst number (intensity), *C*: control and *T*: treated)^59^ was determined. Each experiment included 2 technical repeats (2 feeding cups per compound) and this was repeated for at least three independent biological experiments per compound. Non-parametric *t*-test (Mann-Whitney) was applied (Graphpad Prism 8.3.0) for statistical analysis.

### Endectocide evaluation

Endectocidal activity was determined using SMFA as above but with 2 μM of each compound in cow blood (100 μL) at 37°C for 35 min feeds. Each 350 mL feeding cup contained 30 *An. coluzzii* females (4 h starved, 2-5 days old). Fully or partially fed females were retained for daily monitoring of mortality under standard insectary rearing conditions, for up to 4 days post-treatment. Ivermectin and DMSO were used as positive and negative controls, respectively. Between 3-5 independent replicates were performed per compound. Mean 4-day mortality was statistically evaluated with ANOVA against the negative DMSO control (Graphpad Prism 8.3.0).

### Data analysis and chemical clustering

Drug classes and biological pathways or protein targets were identified for each compound after text and structure searches of PubChem (https://pubchem.ncbi.nlm.nih.gov/), DrugBank (https://www.drugbank.ca/) and SciFinder (https://sso.cas.org). Chemical space analysis was performed with StarDrop v 6.6 (https://www.optibrium.com/stardrop/) based on structure similarities. The launched drug space was generated from the data file available with Stardrop software. The antimalarial drug space was generated using marketed antimalarial drugs and compounds undergoing clinical trials. The connectivity network was constructed by clustering the compounds using the *FragFP* descriptor (Tanimoto similarity index >0.50) in OSIRIS DataWarrior v 5.0.0 (www.openmolecules.org). The network was visualized using Cytoscape v 3.7.2. Supra-hexagonal maps was generated in Rstudio with the RColorBrewer R package.^60, 61^

## Code availability

All computer codes used to analyze the data are available from the corresponding author upon request

## Data availability

The data presented in this study area available from the corresponding author upon request.

## Acknowledgements

We thank the MMV for assembly and supply of the PRB and Didier Leroy and Esperanza Herreros from the MMV for helpful discussions. We thank Dr A. Bennett for prior optimization of the SMFA.

We thank the Medicines for Malaria Venture and South African Technology Innovation Agency (TIA) for funding (Project MMV18/0001). This project was in part supported by the South African Medical Research Council with funds received from the South African Department of Science and Innovation, in partnership with the Medicines for Malaria Venture (KC, LMB, LLK and TLC); and the DST/NRF South African Research Chairs Initiative Grant (LMB UID 84627, LLK UID: 171215294399; KC UID: 64767); and CSIR Parliamentary Grant funding (AT, DM). EAW thanks the Bill and Melinda Gates Foundation for funding (Grant OPP1054480).

## Author contributions

JD, LMB, KC, LLK, GB, EAW and TLC conceptualized the work and supervised the data acquisition. MvdW, JR, DT, NM, SO, AT, PM, SL, BB, EE, NV and LN performed experiments and data analysis. CLM, AvH, AH, GB, DC, LLK and LMB performed additional data analysis. LMB, MvW and JR wrote the manuscript with contributions from all authors.

## Additional Information

Supplementary data files (File S1, S2 and S3) as well as supplementary figures (Fig S1, Fig S2, S3 and Fig S4) are included.

---

**Figure S1:**
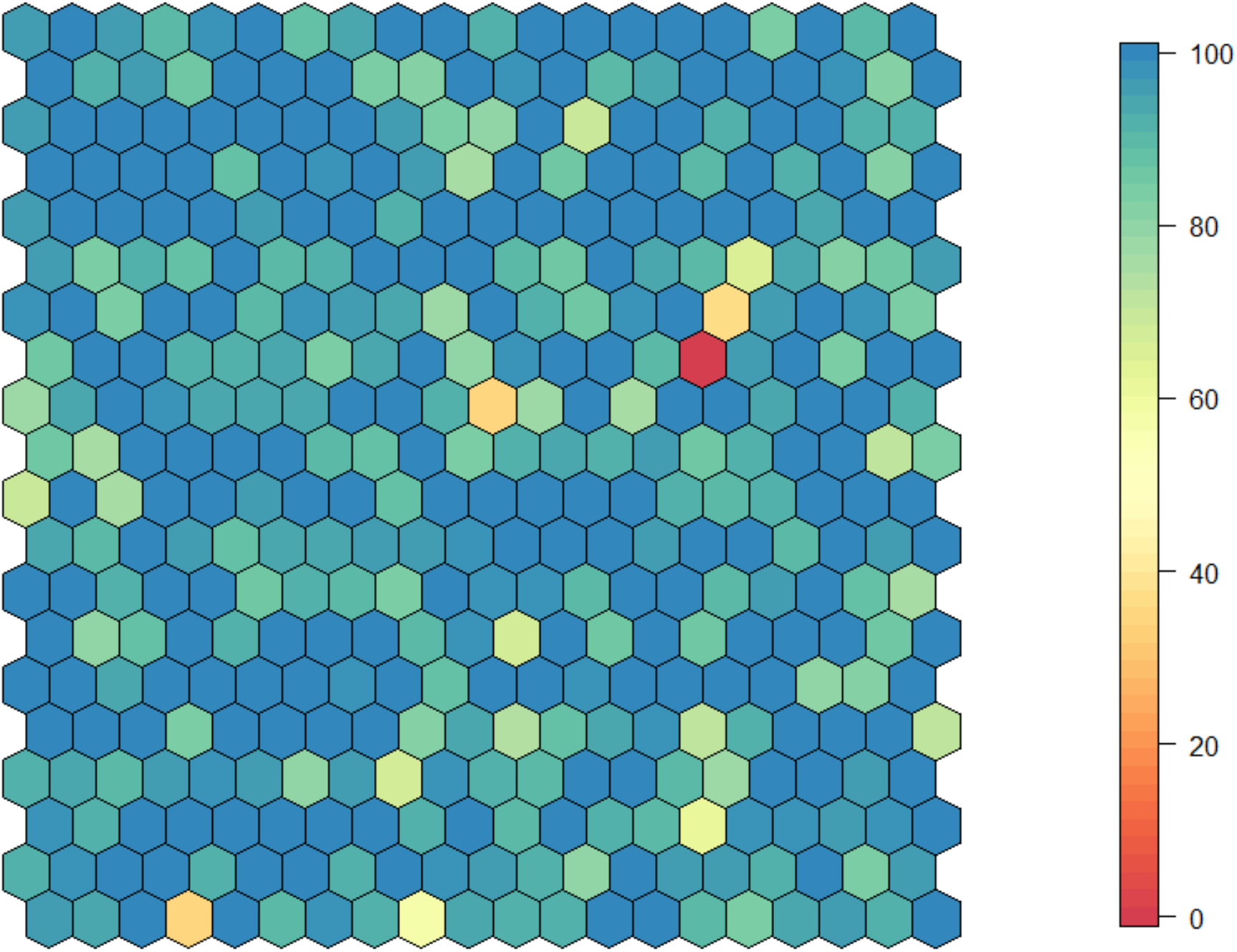
Hexoplot for % inhibition of compounds at 2 uM on CHO cells. Figure supplementary to Figure 2A.

**Figure S2:**
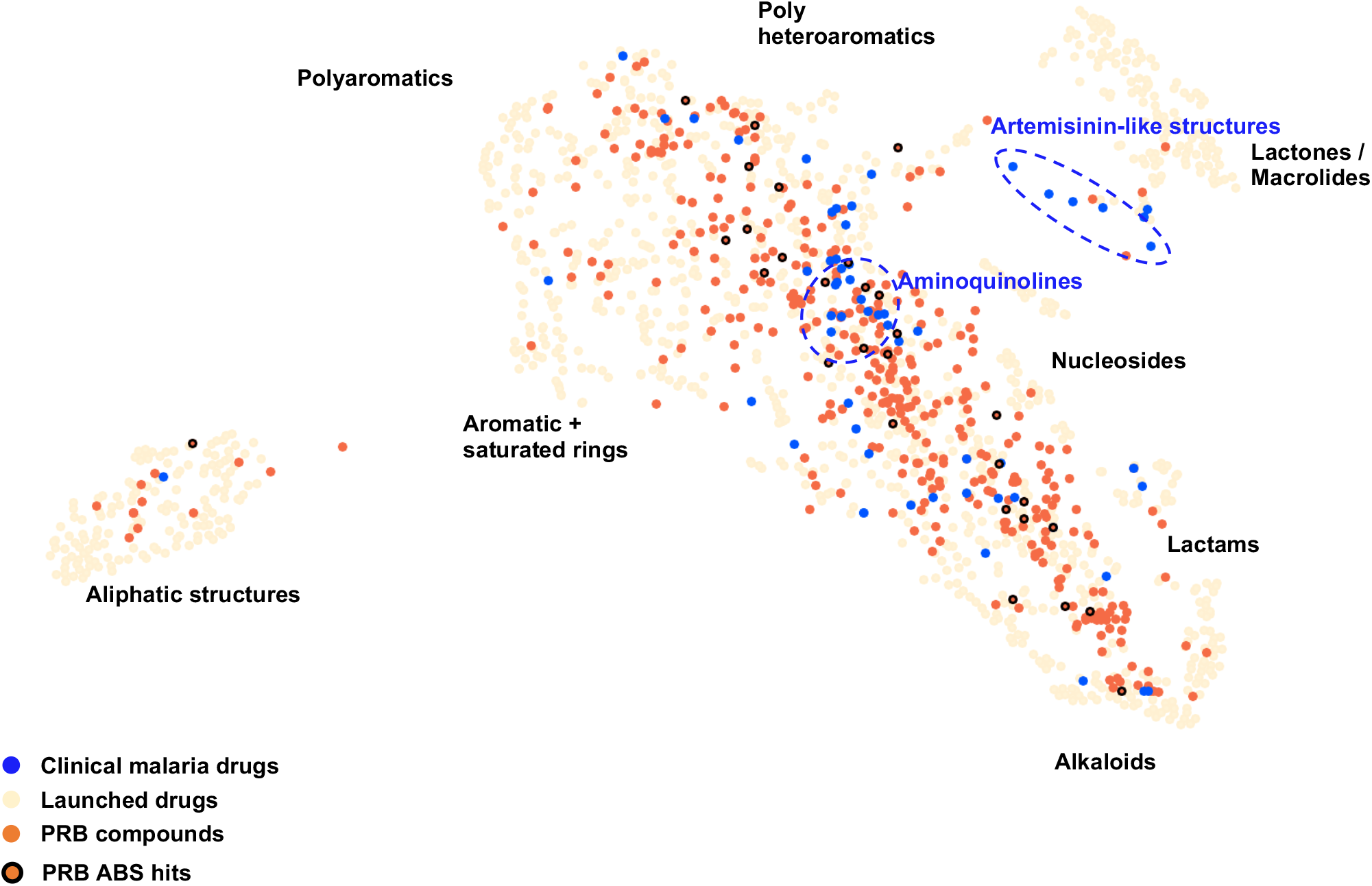
Structural diversity of the MMV PRB compared to Malaria Clinical Drugs. MMV PRB (red dots) and Malaria Clinical Drugs (blue dots) chemical spaces are plotted in the Launched Drugs Space (beige dots).

**Figure S3:**
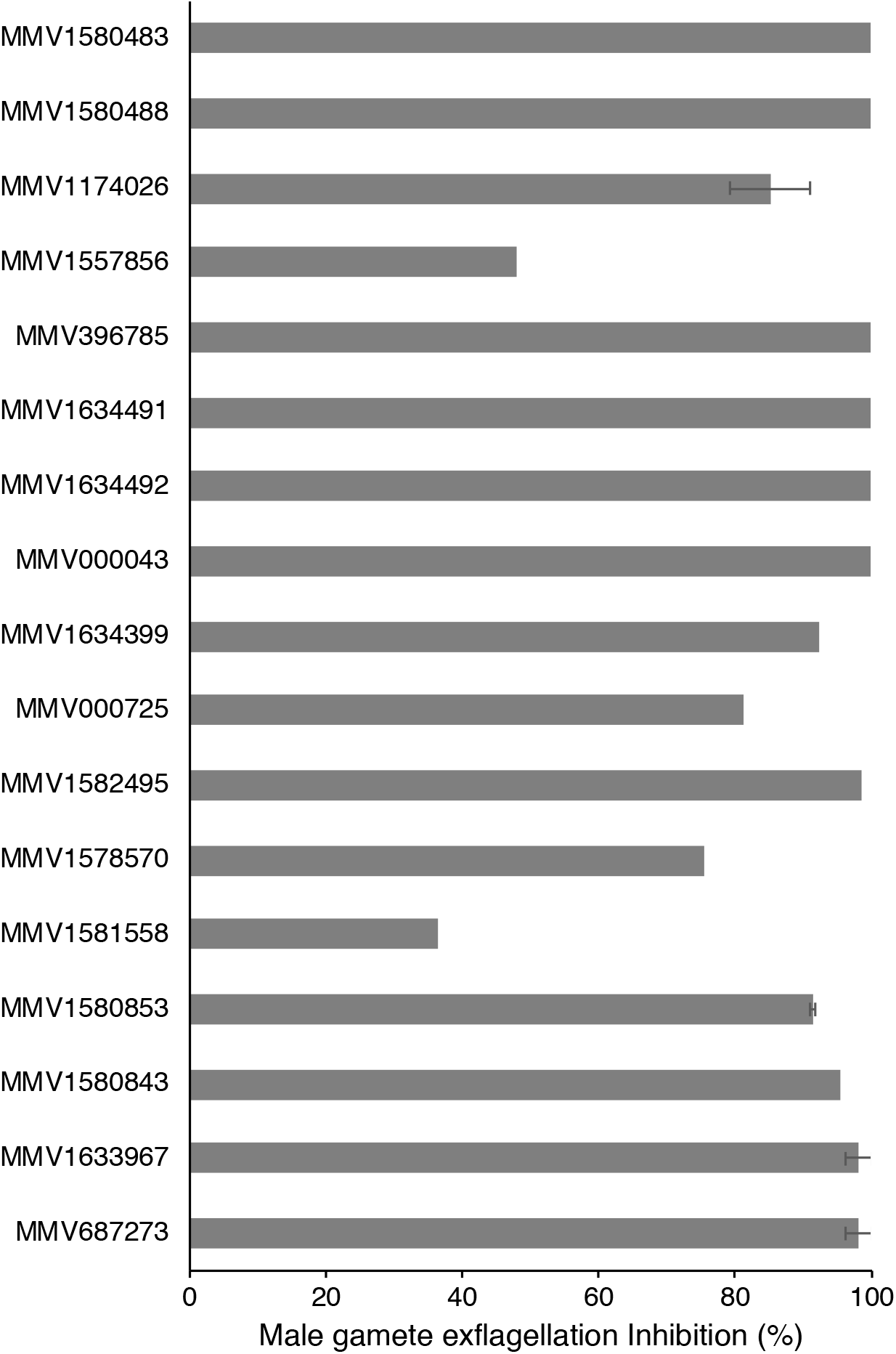
Inhibition of male gamete exflagellation in washout format. Figure supplementary to Figure 5A.

**Figure S4:**
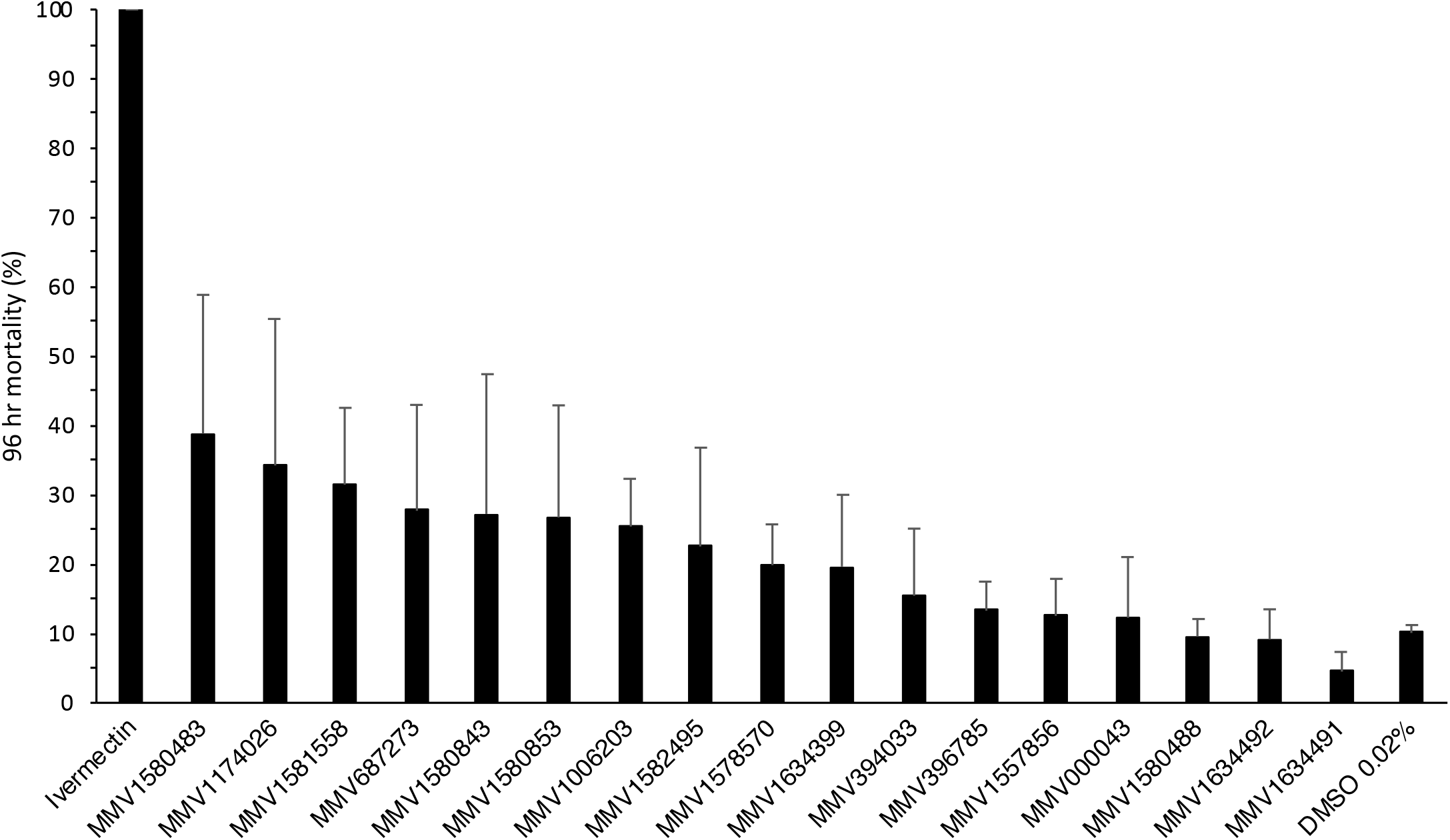
Endectocidal activity of 17 compounds with gametocytocidal activity. Compounds were screened at 2 μM and mortality of *An. coluzzii* (G3) determined after 4 days, compared to Ivermectin (2 μM) as drug control, with DMSO (0.02%) as vehicle control. Data are from ≥2 independent biological repeats, ± S.E.

## References

1. Sinden RE. A biologist’s perspective on malaria vaccine development. Hum Vaccin 6, 3–11 (2010).

2. Josling GA, Llinas M. Sexual development in *Plasmodium* parasites: knowing when it’s time to commit. Nat Rev Microbiol 13, 573–587 (2015).

3. Sinden RE. The cell biology of malaria infection of mosquito: advances and opportunities. Cell Microbiol 17, 451–466 (2015).

4. Reader J, et al. Nowhere to hide: interrogating different metabolic parameters of *Plasmodium falciparum* gametocytes in a transmission blocking drug discovery pipeline towards malaria elimination. Malar J 14, 213 (2015).

5. Plouffe DM, et al. High-Throughput Assay and Discovery of Small Molecules that Interrupt Malaria Transmission. Cell Host Microbe 19, 114–126 (2016).

6. Delves M, et al. The activities of current antimalarial drugs on the life cycle stages of *Plasmodium:* a comparative study with human and rodent parasites. PLoS Med 9, e1001169 (2012).

7. Abraham M, et al. Probing the Open Global Health Chemical Diversity Library for multistage-active starting points for next-generation antimalarials. ACS Infect Dis, (2020).

8. Delves MJ. *Plasmodium* cell biology should inform strategies used in the development of antimalarial transmission-blocking drugs. Future Med Chem 4, 2251–2263 (2012).

9. Delves M, et al. Fueling Open Innovation for Malaria Transmission-Blocking Drugs: Hundreds of Molecules Targeting Early Parasite Mosquito Stages. Front Microbiol 10, 2134 (2019).

10. Miguel-Blanco C, et al. Hundreds of dual-stage antimalarial molecules discovered by a functional gametocyte screen. Nat Commun 8, 15160 (2017).

11. Delves MJ, et al. A high throughput screen for next-generation leads targeting malaria parasite transmission. Nat Commun 9, 3805 (2018).

12. Swann J, et al. High-Throughput Luciferase-Based Assay for the Discovery of Therapeutics That Prevent Malaria. ACS Infect Dis 2, 281–293 (2016).

13. Antonova-Koch Y, et al. Open-source discovery of chemical leads for next-generation chemoprotective antimalarials. Science 362, (2018).

14. Benetatos CA, et al. Birinapant (TL32711), a bivalent SMAC mimetic, targets TRAF2-associated cIAPs, abrogates TNF-induced NF-kappaB activation, and is active in patient-derived xenograft models. Mol Cancer Ther 13, 867–879 (2014).

15. Fernandez-Villa D, Aguilar MR, Rojo L. Folic Acid Antagonists: Antimicrobial and Immunomodulating Mechanisms and Applications. Int J Mol Sci 20, (2019).

16. Frey KM, Viswanathan K, Wright DL, Anderson AC. Prospective screening of novel antibacterial inhibitors of dihydrofolate reductase for mutational resistance. Antimicrob Agents Chemother 56, 3556–3562 (2012).

17. Lamb KM, N GD, Wright DL, Anderson AC. Elucidating features that drive the design of selective antifolates using crystal structures of human dihydrofolate reductase. Biochemistry 52, 7318–7326 (2013).

18. Kirkpatrick JE, Kirkwood KL, Woster PM. Inhibition of the histone demethylase KDM4B leads to activation of KDM1A, attenuates bacterial-induced pro-inflammatory cytokine release, and reduces osteoclastogenesis. Epigenetics 13, 557–572 (2018).

19. Tahlan K, et al. SQ109 targets MmpL3, a membrane transporter of trehalose monomycolate involved in mycolic acid donation to the cell wall core of *Mycobacterium tuberculosis*. Antimicrob Agents Chemother 56, 1797–1809 (2012).

20. Sacksteder KA, Protopopova M, Barry CE, 3rd, Andries K, Nacy CA. Discovery and development of SQ109: a new antitubercular drug with a novel mechanism of action. Future Microbiol 7, 823–837 (2012).

21. Li K, et al. Multitarget drug discovery for tuberculosis and other infectious diseases. J Med Chem 57, 3126–3139 (2014).

22. Ramesh R, et al. Repurposing of a drug scaffold: Identification of novel sila analogues of rimonabant as potent antitubercular agents. Eur J Med Chem 122, 723–730 (2016).

23. Veiga-Santos P, et al. SQ109, a new drug lead for Chagas disease. Antimicrob Agents Chemother 59, 1950–1961 (2015).

24. Torres-Rodriguez JM, et al. In vitro susceptibilities of clinical yeast isolates to the new antifungal eberconazole compared with their susceptibilities to clotrimazole and ketoconazole. Antimicrob Agents Chemother 43, 1258–1259 (1999).

25. Zampieri D, Mamolo MG, Laurini E, Scialino G, Banfi E, Vio L. Antifungal and antimycobacterial activity of 1-(3,5-diaryl-4,5-dihydro-1H-pyrazol-4-yl)-1H-imidazole derivatives. Bioorg Med Chem 16, 4516–4522 (2008).

26. Delves MJ, et al. Male and female *Plasmodium falciparum* mature gametocytes show different responses to antimalarial drugs. Antimicrob Agents Chemother 57, 3268–3274 (2013).

27. Ruecker A, et al. A male and female gametocyte functional viability assay to identify biologically relevant malaria transmission-blocking drugs. Antimicrob Agents Chemother 58, 7292–7302 (2014).

28. Li T, et al. Robust, reproducible, industrialized, standard membrane feeding assay for assessing the transmission blocking activity of vaccines and drugs against *Plasmodium falciparum*. Malar J 14, 150 (2015).

29. Dechering KJ, et al. Modelling mosquito infection at natural parasite densities identifies drugs targeting EF2, PI4K or ATP4 as key candidates for interrupting malaria transmission. Sci Rep 7, 17680 (2017).

30. Colmenarejo G, et al. Predicting transmission blocking potential of anti-malarial compounds in the Mosquito Feeding Assay using *Plasmodium falciparum* Male Gamete Inhibition Assay. Sci Rep 8, 7764 (2018).

31. Vos MW, et al. A semi-automated luminescence based standard membrane feeding assay identifies novel small molecules that inhibit transmission of malaria parasites by mosquitoes. Sci Rep 5, 18704 (2015).

32. Hooft van Huijsduijnen R, et al. Anticancer properties of distinct antimalarial drug classes. PLoS One 8, e82962 (2013).

33. Nzila A, Okombo J, Becker RP, Chilengi R, Lang T, Niehues T. Anticancer agents against malaria: time to revisit? Trends Parasitol 26, 125–129 (2010).

34. Gaillard T, Madamet M, Tsombeng FF, Dormoi J, Pradines B. Antibiotics in malaria therapy: which antibiotics except tetracyclines and macrolides may be used against malaria? Malar J 15, 556 (2016).

35. Pongratz P, Kurth F, Ngoma GM, Basra A, Ramharter M. In vitro activity of antifungal drugs against *Plasmodium falciparum* field isolates. Wien Klin Wochenschr 123 Suppl 1, 26–30 (2011).

36. Burrows JN, et al. New developments in anti-malarial target candidate and product profiles. Malar J 16, 26 (2017).

37. Ebert G, et al. Targeting the Extrinsic Pathway of Hepatocyte Apoptosis Promotes Clearance of *Plasmodium* Liver Infection. Cell Rep 30, 4343–4354 e4344 (2020).

38. Manickam Y, Chaturvedi R, Babbar P, Malhotra N, Jain V, Sharma A. Drug targeting of one or more aminoacyl-tRNA synthetase in the malaria parasite *Plasmodium falciparum*. Drug Discov Today 23, 1233–1240 (2018).

39. Cabrera DG, Horatscheck A, Wilson CR, Basarab G, Eyermann CJ, Chibale K. Plasmodial Kinase Inhibitors: License to Cure? J Med Chem 61, 8061–8077 (2018).

40. Churcher TS, et al. Probability of Transmission of Malaria from Mosquito to Human Is Regulated by Mosquito Parasite Density in Naive and Vaccinated Hosts. PLoS Pathog 13, e1006108 (2017).

41. Burrows J, et al. A discovery and development roadmap for new endectocidal transmission-blocking agents in malaria. Malar J 17, 462 (2018).

42. van Biljon R, et al. Hierarchical transcriptional control regulates *Plasmodium falciparum* sexual differentiation. BMC Genomics 20, 920 (2019).

43. Huy NT, et al. Effect of antifungal azoles on the heme detoxification system of malarial parasite. J Biochem 131, 437–444 (2002).

44. Nagaraj VA, et al. Malaria parasite-synthesized heme is essential in the mosquito and liver stages and complements host heme in the blood stages of infection. PLoS Pathog 9, e1003522 (2013).

45. Matthews KA, et al. Disruption of the *Plasmodium falciparum* life cycle through transcriptional reprogramming by inhibitors of Jumonji demethylases. ACS Infect Dis, (2020).

46. Zhang B, et al. Crystal Structures of Membrane Transporter MmpL3, an Anti-TB Drug Target. Cell 176, 636–648 e613 (2019).

47. MacRae JI, et al. Mitochondrial metabolism of sexual and asexual blood stages of the malaria parasite *Plasmodium falciparum*. BMC Biol 11, 67 (2013).

48. Gulati S, et al. Profiling the Essential Nature of Lipid Metabolism in Asexual Blood and Gametocyte Stages of Plasmodium falciparum. Cell Host Microbe 18, 371–381 (2015).

49. G. Costa MG, M. Eldering, R.L. Lindquist, A.E. Hauser, R. Sauerwein, C. Goosmann, V. Brinkmann, View ORCID Profile E.A. Levashina. Non-competitive resource exploitation within-mosquito shapes evolution of malaria virulence.

50. Adjalley SH, Johnston GL, Li T, Eastman RT, Ekland EH. Quantitative assessment of *Plasmodium falciparum* sexual development reveals potent transmission-blocking activity by methylene blue. Proceedings of the National Academy of Sciences of the United States of America 108, 1214 – 1223 (2011).

51. Wu CP, van Schalkwyk DA, Taylor D, Smith PJ, Chibale K. Reversal of chloroquine resistance in *Plasmodium falciparum* by 9H-xanthene derivatives. Int J Antimicrob Agents 26, 170–175 (2005).

52. Makler MT, et al. Parasite lactate dehydrogenase as an assay for *Plasmodium falciparum* drug sensitivity. Am J Trop Med Hyg 48, 739–741 (1993).

53. Paquet T, et al. Antimalarial efficacy of MMV390048, an inhibitor of *Plasmodium* phosphatidylinositol 4-kinase. Sci Transl Med 9, (2017).

54. Coetzee N, von Gruning H, Opperman D, van der Watt M, Reader J, Birkholtz LM. Epigenetic inhibitors target multiple stages of *Plasmodium falciparum* parasites. Sci Rep 10, 2355 (2020).

55. Rubinstein LV, et al. Comparison of in vitro anticancer-drug-screening data generated with a tetrazolium assay versus a protein assay against a diverse panel of human tumor cell lines. J Natl Cancer Inst 82, 1113–1118 (1990).

56. Coetzee M, Hunt RH, Wilkerson R, Della Torre A, Coulibaly MB, Besansky NJ. *Anopheles coluzzii* and *Anopheles amharicus,* new members of the Anopheles gambiae complex. Zootaxa 3619, 246–274 (2013).

57. Fanello C, Santolamazza F, della Torre A. Simultaneous identification of species and molecular forms of the *Anopheles gambiae* complex by PCR-RFLP. Med Vet Entomol 16, 461–464 (2002).

58. Hunt RH, Brooke BD, Pillay C, Koekemoer LL, Coetzee M. Laboratory selection for and characteristics of pyrethroid resistance in the malaria vector *Anopheles funestus*. Med Vet Entomol 19, 271–275 (2005).

59. Miura K, et al. Transmission-blocking activity is determined by transmission-reducing activity and number of control oocysts in *Plasmodium falciparum* standard membrane-feeding assay. Vaccine 34, 4145–4151 (2016).

60. Nychka D, Furrer R, Paige J, Sain S. fields: Tools for spatial data. (2017).

61. Neuwirth E. RColorBrewer: ColorBrewer Palettes. (2014).

